# Inflammation plays a critical role in damage to the bronchiolar epithelium induced by *Trueperella pyogenes in vitro* and *in vivo*

**DOI:** 10.1101/2023.07.24.550341

**Authors:** Lei Qin, Fandan Meng, Haijuan He, Siqi Li, Hongliang Zhang, Yuan Sun, Wenlong Zhang, Tongqing An, Xuehui Cai, Shujie Wang

## Abstract

*Trueperella pyogenes* can cause severe pulmonary disease in swine, but the mechanism of pathogenesis is not well defined. *T. pyogenes-*induced damage to porcine bronchial epithelial cells (PBECs), porcine precision-cut lung slices (PCLS) and respiratory epithelium of mice remains unknown. In this study, we used *T. pyogenes* 20121 to infect PBECs in air-liquid interface conditions and porcine PCLS. *T. pyogenes* could adhere to, colonize and induce cytotoxic effect on PBECs and the luminal surface of bronchi in PCLS, which damaged the bronchiolar epithelium. Moreover, bronchiolar epithelial cells showed extensive degeneration in infected mice lungs. Furthermore, western blot showed the NOD-like receptor (NLR)/ C-terminal caspase recruitment domain (ASC)/caspase-1 axis and nuclear factor-kappa B (NF-κB) pathway were involved in inflammation in PCLS and lungs of mice, which also confirms PCLS provide a platform to analyze pulmonary immune response. Meanwhile, the levels of p-c-Jun N-terminal kinase (JNK), p-extracellular signal-regulated kinase (ERK) and p-protein kinase B (AKT) were increased significantly, which indicated the mitogen-activated protein kinase (MAPK) and Akt pathways were also involved of inflammation in *T. pyogenes-*infected mice. In addition, we used *T. pyogenes* 20121 to infect tumour necrosis factor alpha (TNF-α)^-/-^ mice, the results indicated apoptosis and injury in respiratory epithelium of infected TNF-α^-/-^ mice were alleviated. Thus, pro-inflammatory cytokine TNF-α played a role in apoptosis and respiratory epithelium injury of mice lungs. Collectively, our study provides an insight into the inflammatory injury induced by *T. pyogenes*, and suggests that blocking NLR or TNF-α may be a potential therapeutic strategy against *T. pyogenes* infection.

## Introduction

*T. pyogenes* is frequently isolated from pyogenic disease conditions in both domestic and wild animals worldwide, but is rare in companion animals and humans. *T. pyogenes* also plays an important role in secondary infection and co-infection in domestic animals (1, 2). In swine, *T. pyogenes* is a common opportunistic pathogen found in pneumonia, endocarditis, pleuritis and organ abscesses (3, 4), and it is an emerging clinical, epidemiological and economic problem on pig farms (5). Although pneumonia in swine with airway inflammation caused by *T. pyogenes* is a frequent occurrence (6), the mechanisms of pathogenesis remain poorly understood.

The primary route of infection for many respiratory pathogens is the airway, which is lined by a layer of epithelial cells that form a primary barrier (7). Thus, well-differentiated porcine bronchial epithelial cells (PBECs) in air-liquid interface (ALI) conditions, which provide a close *in vitro* model of the airway epithelium (8, 9), have been used to study bacterial infections (10, 11). Precision-cut lung slices (PCLS) as ex-vivo lung culture can mimic the immediate and long-term functional responses of the respiratory tract and lung (12–14), which allows studies on inflammatory responses induced by respiratory pathogens (15, 16). Moreover, the 3D respiratory organotypic tissue reflects the natural microanatomy and microenvironment of the respiratory system (14), permitting a reduction in the number of laboratory animals used (33, 29).

Little is known about inflammatory response and mechanism of bronchial damage of *T. pyogenes* infections *in vitro* or *in vivo*, though we recently used PCLS to study the pathogenicity of a novel isolate (17). In the present study, ALI cultures, PCLS and mice were used to analyze the adherence and colonization of *T. pyogenes* to porcine airway epithelial cells, seeking to further elucidate how the bacterium damages the respiratory tract.

## Materials and Methods

### Bacterial strain and growth conditions

The virulent strain *T. pyogenes* 20121 was isolated from the lungs of sick pigs from a pig farm as previously described (17) and stored in our lab. *T. pyogenes* 20121 was grown in tryptic soy broth (TSB; Difco, Loveton Circle Sparks, MD, USA) supplemented with 5% fetal bovine serum (FBS; CLARK, USA) or on Columbia-based blood agar media (ThermoFisher Scientific, China) at 37 °C. For preparation of cryo-conserved bacterial stocks, *T. pyogenes* was grown in TSB medium until late exponential growth phase (OD_600nm_ = 0.5). Bacteria were harvested by centrifugation (3000 g for 5 min at 4 °C), washed once with phosphate-buffered saline (PBS) and re-suspended in TSB medium containing 50% (v/v) glycerol. Aliquots were immediately shock frozen in liquid nitrogen and stored at −80 °C until use.

### Infection of well-differentiated PBECs by *T. pyogenes*

Porcine lungs were obtained from specific pathogen free (SPF) pigs, and primary PBECs were harvested from the bronchi as previously described (11). Briefly, PBECs were scraped carefully from the luminal surface of the bronchus without disturbing the mucosal surface integrity and cultured in bronchial epithelial cell growth medium (BEBM; Lonza, Belgium) supplemented with antibiotics. When PBECs reached about 80% confluence, they were seeded on 0.4 μm pore size transwell filters with polycarbonate membranes (Corning Costar, USA). Cells were then cultured at 37 °C with 5% CO_2_ with ALI medium as described previously (18), which is mixture of BEBM and DMEM at 1:1 supplemented with antibiotics. At 3 days post-seeding, cultures were maintained under ALI conditions for an additional 4 weeks at 37 °C with 5% CO_2_ for epithelial cell differentiation, changing the culture medium every two days.

Growth characteristics of *T. pyogenes* 20121 was determined on PBECs that cultured under air-liquid interface condition for 4 weeks, and all treatments were repeated at least three times. Transwell filters were washed 3 times with PBS and cultured in fresh medium without antibiotics for 24 h prior to infection. *T. pyogenes* 20121 was inoculated into the apical compartment with approximately 8×10^5^ CFU per filter, and the supernatant from apical compartment was collected for a cytotoxicity assay after 4 h at 37 °C with 5% CO_2_. Both apical and basolateral compartments were washed 3 times with PBS to remove any non-adherent bacteria, and maintained under ALI condition. Cytotoxicity assays and immunofluorescence on PBECs were performed at 4 h post-inoculation (hpi), 24 hpi, 48 hpi and 72 hpi.

### Infection of *T. pyogenes* on porcine PCLS

The cranial, middle and intermediate lobes were harvested from fresh lung tissue of three sacrificed SPF pigs. Each lobe was gently filled with 1.5% (w/v) warming low-melting agarose (Promega, USA) along the bronchus as described previously (19). The lobes were stamped out with an 8-mm tissue coring tool, and after the agar was solidified, slices were prepared on a Krumdieck tissue slicer (model MD6000-01; TSE Systems, USA) with 6-10 slices per minute. Next, the PCLS were carefully transferred into 24-well plates and maintained with 1 mL fresh RPMI medium 1640 supplemented with antibiotics (including 80 μg/ml kanamycin, 10 μg/ml enrofloxacin, 5 μg/ml levofloxacin, 2.5 μg/ml amphotericin, 50 μg/ml ciprofloxacin), and the medium was refreshed per hour to remove the agarose and repeat for 3 times, then cultured overnight. PCLS containing bronchia and showing 100% ciliary activity were selected by light microscopy (EVOS FL Auto; Thermo Fisher Scientific, USA) for subsequent experiments. RPMI medium 1640 without antibiotic or antimycotic was added 24 h prior to bacterial infection studies. PCLS were washed 3 times with PBS, then inoculated with *T. pyogenes* at 8×10^4^ (3 slices) or 8×10^5^ CFU (3 slices) per slice at 37 °C, and the control group (mock-infected slices, 6 slices total) was cultured normally under the same conditions. The experiment was repeated three times (a pig per time, 3 pigs total). At 4 hpi, slices were washed three times with PBS to remove non-attached bacteria, and 1 mL fresh RPMI medium 1640 (Gibco, Beijing, China) was added for further cultivation. The areas of bronchial cavity were measured and calculated by ImageJ/Fiji software, and bronchial contraction percentage (BCP) presented the results using the following formula, BCP = [reduced bronchial cavity area / initial bronchial cavity area] × 100%. The supernatant was collected at 4 hpi, 24 hpi and 72 hpi for cytotoxicity assays, and PCLS were homogenized in PBS and then serially diluted and plated onto TSA for enumeration of bacteria.

### Cytotoxicity assay

The release of lactate dehydrogenase (LDH, Promega, USA) into the culture medium was quantitated using Cyto Tox 96® Non-Radioactive Cytotoxicity Assay (Promega, Madison, WI, USA) as described previously (20). To determine the cytotoxic effect caused by *T. pyogenes* 20121, 50 μL of supernatant from infected or mock-infected PBECs or PCLS at each time point was mixed with an equal volume of substrate mix reagent in the dark. The detected supernatant in PBECs were obtained by washing the upper chamber of transwell for 20 minutes using culture medium on the shaker at 24, 48, 72 hpi. After 30 min, stop solution was added and absorbance signal was measured at 490 nm. The results were compared with the control group and presented as 100% cytotoxicity. Three parallel PBECs or PCLS (from the same pig) in different infected groups were collected at each detection time, and the infected experiments were repeated three time (a pig per time, 3 pigs total). The cytotoxicity assay in PBECs or PCLS were replicated three times.

### Animal experiments

All animal experiments were performed in accordance with the Guide for the Care and Use of Laboratory Animals of the Ministry of Science and Technology of the People’s Republic of China. Mouse infection experiments (approval number 210119-02) were carried out in the animal biosafety level 2 facilities under supervision of the Committee on the Ethics of Animal Experiments of the Harbin Veterinary Research Institute of the Chinese Academy of Agricultural Sciences (CAAS) and the Animal Ethics Committee of Heilongjiang Province, China. 7-week-old female C57BL/6 wild type (WT) mice (Changsheng Biotechnology, Liaoning, China) were randomly divided into 2 groups. Uninfected controls (n=12) received 0.2 mL TSB intraperitoneally (i.p.), and the rest (n=20) were challenged i.p. with 0.2 mL TSB containing 2 × 10^6^ CFU *T. pyogenes* 20121. Lung and blood samples from infected mice (n=5) and control (n=3) were aseptically harvested after 1, 2, 4 and 7 days post-infection (dpi) for histopathological analysis and cytokine detection. Bacterial load in lungs and blood were monitored as previously described (21), briefly, anticoagulated blood and lung samples were serially diluted, and plated on blood agar medium for 36 h, and the bacteria were quantified by colony counting. In a separate experiment, 16 female TNF-α^-/-^ C57BL/6N mice were purchased from Cyagen Biosciences (Santa Clara, USA), and at 6-7 weeks-of-age were randomly divided into 2 groups. Uninfected controls (n=6) received 0.2 mL TSB i.p., and the other group was inoculated i.p. with *T. pyogenes* 20121 as above. 5 infected TNF-α^-/-^ mice and 3 uninfected TNF-α^-/-^ mice were humanely euthanized at 1 and 2 dpi, Lung and blood samples were harvested, and lung tissue samples were fixed in 3.7% formaldehyde (Amresco, Fountain Parkway, USA) for further histopathological analysis. Lung samples were sectioned (8-μm-thick slices) on a cryostat and used for double-immunofluorescence staining.

### Cryosections and immunofluorescence analysis

PCLS were washed 3 times using PBS to remove unattached bacteria, embedded on filter paper by tissue freezing medium (Sigma-Aldrich, USA) and quickly frozen in liquid nitrogen, then stored at −80 °C. 10-μm-thick cryo-slices were produced by a cryostat (Thermo, USA) and stored at −20 °C. The slices were dried at room temperature (RT) prior to immunofluorescence analysis.

PBECs and cryosection samples were fixed with 3.7% paraformaldehyde (Amresco, Fountain Parkway, USA) for 30 min, followed by 0.1 M glycine treatment for 20 min at RT. After three washing steps with PBS, samples were permeabilized with 0.2% Triton X-100 (Sigma-Aldrich, USA) for 20 min at RT. 1% (v/v) bovine serum albumin (BSA; biofroxx, Germany) was used to block nonspecific reactions for 30 min at RT. The primary antibody for detection of *T. pyogenes* was polyclonal mouse antiserum (1: 300, made in our lab), and ciliated cells were stained using a Cy3-labeled anti-β-tubulin monoclonal antibody (1:300; Sigma-Aldrich, USA). Alexa Fluor^®^ 488-labeled goat-anti-mouse IgG (H+L) (1:1000; Thermo Fisher Scientific, USA) was used as secondary antibody; all antibodies were diluted in 1% BSA. Nuclei were stained with DAPI (4′,6-diamidino-2-phenylindole; Cell Signaling Technology, USA), and finally samples were mounted with ProLong^®^ Gold Antifade Reagent (Thermo Fisher Scientific, USA) and stored at 4 °C with light protection. Slides were examined on a confocal laser scanning microscopy with fast Airyscan (LSM980-ZEISS, Germany), and Z-stack images were acquired containing 0.22 μm per plane. Maximum intensity projections were calculated for display purposes and adjusted for brightness and contrast using ZEN 2.3 blue software.

Lung samples collected at necropsy were sectioned (7 μm-thick) and used for apoptosis detection. To confirm the cells of lungs that underwent apoptosis, lung sections were stained with rabbit anti-mouse caspase-3 antibody (1:500, Cell Signaling, MA, USA) and Alexa Fluor^TM^ 488-conjugated goat anti-rabbit antibody (1:1000, Sigma, Missouri, USA); Apoptosis also was detected using a terminal deoxynucleotidyl transferase (TdT)-mediated deoxyuridine triphosphate (dUTP)-biotin nick end-labeling (TUNEL) assay with Cell Death Detection Kit (Roche, Germany). Nuclei were stained with 4-6-diamidino-2-phenylindole (DAPI, Sigma).

### Western blot analysis

Lung samples from the PCLS and mice were lysed in RIPA buffer containing the protease inhibitors PMSF (Solarbio, China) and Complete Protease Inhibitor (EDTA-free; Merck-Millipore, Germany). Protein concentration was quantified using a BCA Protein Assay Kit (Beyotime Institute of Biotechnology, China). Equal amounts of protein were loaded and separated by electrophoresis on 12% SDS-PAGE gels and subsequently transferred to polyvinylidene fluoride (PVDF; Merck-Millipore, Germany). After blocking with 10% skim milk for 1 h, the membranes were incubated at room temperature for 1 h with the antibodies about the inflammasome, including the rabbit anti-NLRP1, NLRC4, gasdermin D (GSDMD), GSDMD-C (1:1,000, abcam), IL-1β, IL-4, IL-18, matrix metalloproteinase 9 (MMP9), macrophage migration inhibitory factor (MIF) (1:500, ABclonal), NLRP3, caspase-1 p20 and the mouse anti-ASC (1:1,000, these three antibodies were kindly provided by Professor Changjiang Weng of Harbin Veterinary Research Institute of Chinese Academy of Agricultural Sciences (22). The rabbit anti-AKT1/2/3 and phosphor-AKT1/2/3, p38 and phospho-p38, ERK1/2 and phospho-ERK1/2, JNK1/2/3 and phospho-JNK1/2/3, NF-κB p65 and phospho-NF-κB p65 (1:1,000, abcam) were prepared for the tests of MAPK, Akt and NF-κB signaling pathways. Furthermore, antibodies related to apoptosis, including the rabbit anti-Caspase-8, Caspase-9, Caspase-3 and apoptosis inducing factor (AIF) (1:1,000, abcam) were also used in this process. Expression of all the target proteins were normalized to that of the internal control rabbit/mouse antibodies against β-actin or GADPH (1:50,000, ABclonal). Relevant DyLight™ 800-labeled goat anti-rabbit/-mouse IgG (H+L) secondary antibodies were applied as needed (SeraCare, KPL Antibodies & Conjugates, USA). The relative integrated density of the target protein to the internal control was quantified using Image J v1.8.0 software (Wayne Rasband, National institutes of Health, USA) and then the relative quantitative comparison was shown after the normalization of the relative integrated density of the control group.

### Detection of cytokine production in PCLS or mice

To quantify the cytokines induced by *T. pyogenes* infection, supernatants from infected PCLS were collected at 4 hpi, 24 hpi and 72 hpi, and the quantity of interleukin (IL)-4, IL-10 and chemokine CXCL8 was determined by ELISA according to the manufactureŕs instructions (USCN Life Sciences, China). The quantity of IL-1β, IL-6, IL-10, TNF-α and IFN-γ was also determined in serum samples collected from infected mice at 1, 2, 4 and 7 dpi by commercial ELISA according to the manufacturer’s instructions (Cusabio, China).

### Statistical analyses

All experiments were performed at least three times, and results are expressed as the mean ± standard deviation (SD). Statistical analysis of the results was performed with one-or two-way analysis of variance (ANOVA) using GraphPad Prism software version 9.00 (GraphPad, San Diego, CA, USA). A *P* value < .05 was considered statistically significant.

## Results

### *T. pyogenes* infection induces epithelium damage on well-differentiated PBECs

The infection of well-differentiated PBECs by *T. pyogenes* was first analyzed under ALI conditions. *T. pyogenes* induced cytotoxic effects as early as 4 h post-infection in comparison to mock-infected PBECs (Fig. 1A). No significant cytotoxic effect was detected at 24 hpi in comparison to mock-infected PBECs, but cytotoxic effect was significantly enhanced at 48 hpi. Attachment of *T. pyogenes* to epithelial cells under ALI condition was detected by confocal microscopy (Fig. 1B). Tiny microcolonies of *T. pyogenes* were observed adhering to the cilia of ciliated cell at 4 hpi. The size of the microcolonies increased by 24 hpi, indicating *T. pyogenes* was able to proliferate on the epithelial cells under ALI condition. Notably, *T. pyogenes* caused severe damage to the epithelial cells and the cilia after infection, and epithelial integrity was lost by 72 hpi. Importantly, *T. pyogenes* crossed the epithelial cell layer and reached the membrane of the transwell filter.

**Figure 1.**
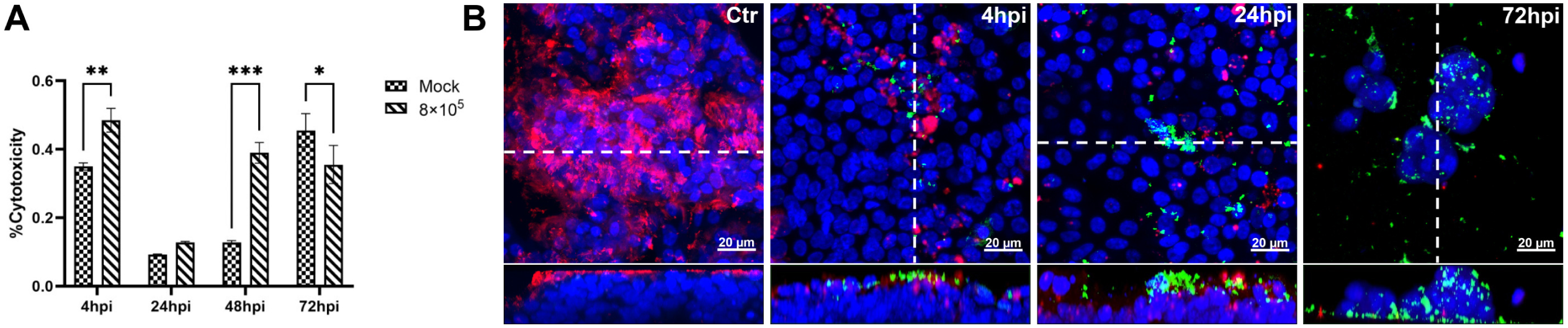
Cytotoxicity and *T. pyogenes*-induced damage to well-differentiated porcine bronchial epithelial cells. PBECs were apically infected with approximately 8×10^5^ CFU of *T. pyogenes* and washed thoroughly to remove non-adherent bacteria after 4 h and further incubated under ALI conditions. **(A)** Cytotoxic effects of *T. pyogenes* on porcine bronchial epithelial cells (PBECs) grown under air-liquid interface (ALI) conditions were quantified by a standard LDH-release assay. Results are expressed as % cytotoxicity compared to 100% killed cells condition and expressed as mean ± SD; ***: *P* < .001, determined using one-way ANOVA and Tukeýs multiple comparison test. Experiments were performed three times. **(B)** Immunostaining was performed to visualize cilia (red) and *T. pyogenes* (green), and nuclei were stained by DAPI (blue). Bars represent 20 μm in horizontal sections for upper images, and lower images are the orthogonal views of Z-stacks (white dotted line) as shown in YZ section at 4 hpi and 72 hpi or XZ section in Control (Ctr) and 24 hpi.

### Adhesion and colonization of *T. pyogenes* on the luminal surface of bronchi in porcine PCLS

To investigate the adhesion and colonization characteristics, porcine PCLS were infected or mock-infected by *T. pyogenes* 20121. Bronchoconstriction was induced by bacterial infection in a dose-dependent manner (Fig. 2A). At 8×10^5^ CFU/slice, *T. pyogenes* caused notable bronchoconstriction (BCP: 94.46%, 87.18%, 84.91%, respectively), while slighter constriction (BCP: 62.69%, 70.09%, 75.02%, respectively) was seen with 8×10^4^ CFU/slice at 4, 24 and 72 hpi. *T. pyogenes* adhered in microcolonies to the luminal surface of bronchi that was presented by the ciliated epithelial cell layer (Fig. 2A). Proliferation of *T. pyogenes* was not affected by bronchoconstriction, with colonizing bacteria still able to proliferate in the constricted area of bronchioles, as indicated by the arrow (Fig. 2A, 24 hpi). Infection with 8×10^5^ CFU of *T. pyogenes* also caused more severe injury to the ciliated epithelial layer than the lower dose at 72 hpi (Fig. 2A). Meanwhile, the CFU counts in the PCLS slice lysate also showed much higher numbers at 24 and 72 hpi than at 4 hpi (Fig. 2B), confirming the proliferation of *T. pyogenes*. Additionally, the 8×10^5^ CFU-infected group had more bacteria attached (>10^5^ CFU) than the 8×10^4^ CFU infected group (10^4^ CFU) at 24 hpi, whereas no significant difference was detected at 72 hpi.

**Figure 2.**
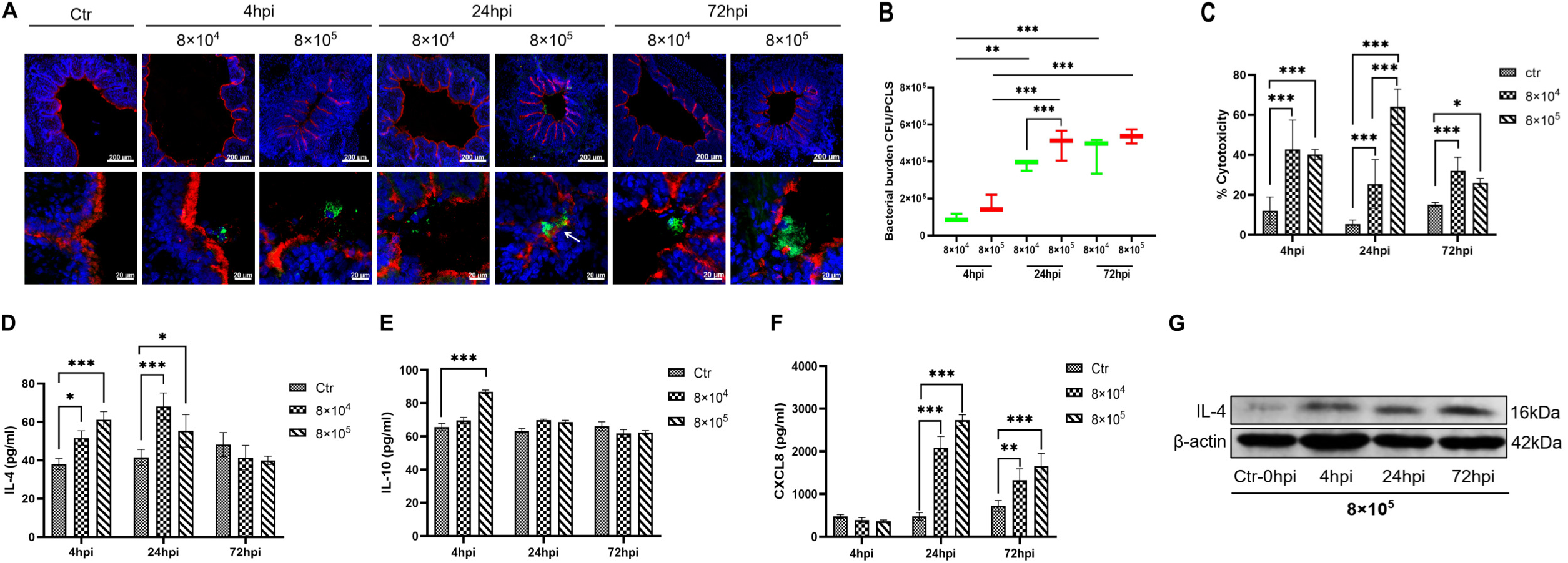
Interaction between *T. pyogenes* with porcine precision-cut lung slices. Precision-cut lung slices (PCLS) were infected with 8×10^4^ or 8×10^5^ CFU/well of *T. pyogenes* for 4 h and washed thoroughly to remove non-adherent bacteria. Cells were fixed for cryosections at 4, 24 and 72 hpi and used for immunofluorescence staining. **(A)** *T. pyogenes* is shown in green, cilia (β-tubulin) in red, and nuclei (DAPI) in blue. **(B)** Bacterial burden of *T. pyogenes* on PCLS at 4, 24, 72 hpi. **(C)** *T. pyogenes*-induced cytotoxicity on PCLS, expressed as % cytotoxicity compared to 100% killed cells condition. **(D)** IL-4 and **(E)** IL-10 levels in PCLS supernatant after incubation with *T. pyogenes.* **(F)** ELISA was used to determine the level of chemokine CXCL8 in culture supernatant. **(G)** Western blot analysis was used to determine the amount of IL-4 in the infected PCLS tissue at 4, 24 and 72hpi. The PCLS from 3 pigs and experiments were performed three times; *: *P* < .05, **: *P* < .01, ***: *P* < .001.

As a measure of cytotoxicity, the amount of LDH released from infected PCLS was determined. *T. pyogenes* induced significant cytotoxic effect on PCLS at 4 and 24 hpi, and the high-dose group caused greater cytotoxicity than the low-dose group (Fig. 2C). The low-dose infection group caused slightly higher cytotoxicity than the high-dose group at 72 hpi, which may be related to a decrease in the number of ciliated epithelial cells due to the more serious damage caused by the high-dose infection.

### Secretion of inflammatory cytokines into the supernatant of infected porcine PCLS

To understand the immune response triggered by *T. pyogenes* in PCLS, the amount of cytokines released into supernatant were measured. Anti-inflammatory cytokines IL-4 and IL-10 in supernatant (Fig. 2D and E) were secreted at 4 hpi, which was earlier than secretion of the chemokine CXCL8 (Fig. 2F). The level of IL-4 in supernatant of both infected groups was significantly higher (*P* < .001) than mock infection at 4 and 24 hpi, and IL-10 increased significantly by 4 hpi only in the high-dose group. Western blot analysis also confirmed an upregulation of IL-4 in PCLS tissue infected with 8×10^5^ CFU at 4, 24 and 72 hpi (Fig. 2G). However, chemokine CXCL8 was notably upregulated in supernatant of *T. pyogenes*-infected PCLS by 24 and 72 hpi compared with mock infection.

### *T. pyogenes* activated the NLRP3 and NF-κB pathways in porcine PCLS

To investigate the effect of the NLRP3 inflammasome on the *T. pyogenes*-induced inflammatory response, PCLS were collected for western blot analysis. *T. pyogenes* infection activated NLRP3 inflammasome, with NLRP3 expression enhanced significantly from 4 hpi to 72 hpi (Fig. 3A), and ASC expression was also enhanced from 4 hpi to 72 hpi compared to control, although it also suggests a drop from 4 h to 24h (Fig. 3B). Meanwhile, pro-caspase-1 was activated and cleaved, with caspase-1 (p20) enhanced significantly at 24 and 72 hpi (Fig. 3C). Furthermore, inflammatory cytokines IL-1β and IL-18 expression were upregulated from 4 to 72 hpi compared to control, but a slight drop in 72 hpi compared to 24 hpi (Fig. 3D and E). In addition, NF-κB and two important factors (MMP9 and MIF) that participate in inflammation were analyzed. Expression of MMP9 increased significantly from 4 to 72 hpi (Fig. 3F) and expression of MIF increased strikingly at 24 and 72 hpi (Fig. 3G) compared to control. Expression of NF-κB increased substantially at 24 hpi and 72 hpi compared to control. These results suggest that the NLRP3 inflammasome and NF-κB pathways are both activated in PCLS infected with *T. pyogenes*.

**Figure 3.**
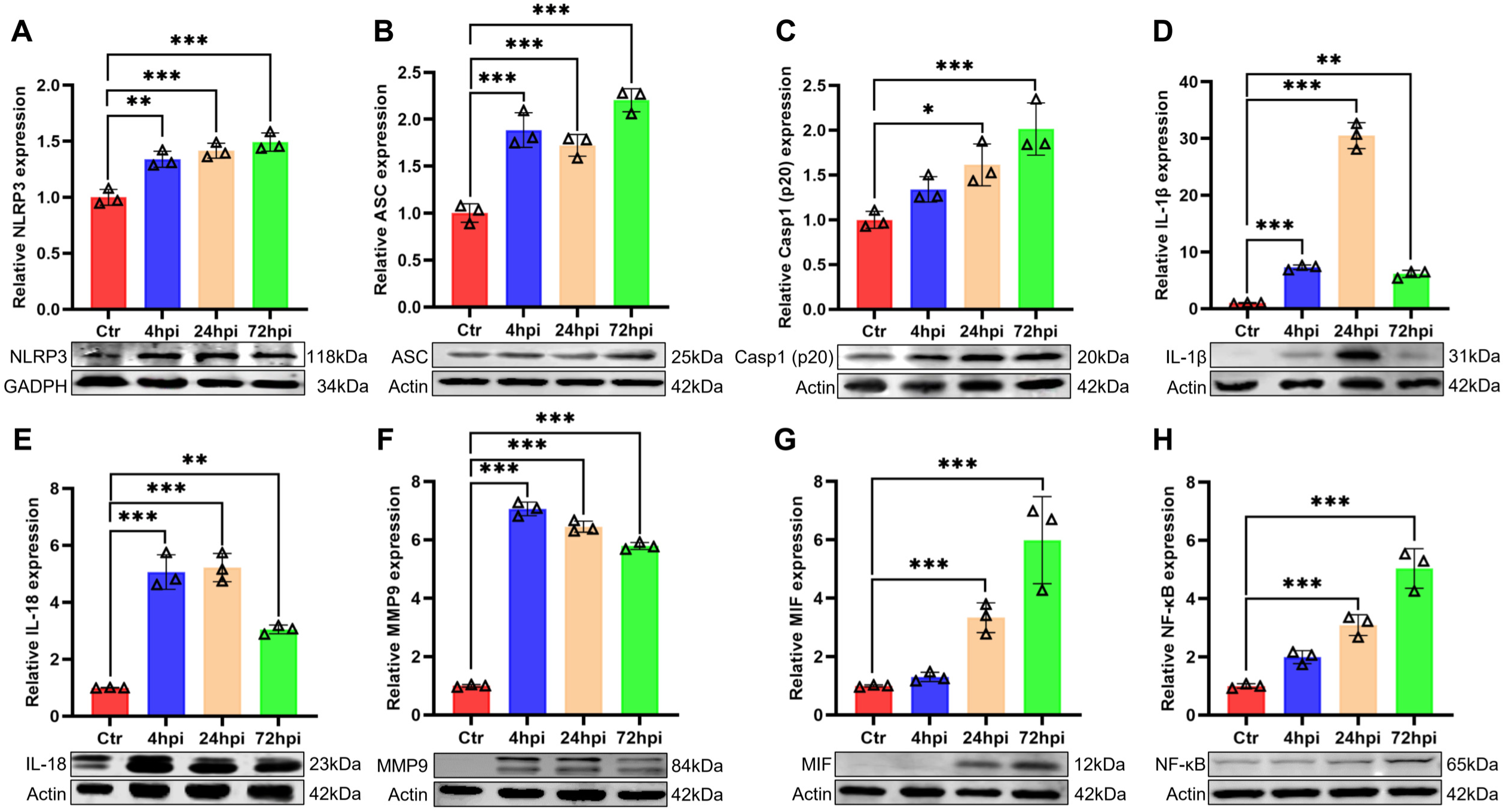
The NLRP3 and NF-κB pathways are activated by *T. pyogenes* in PCLS. Precision-cut lung slices (PCLS) were challenged with 8×10^5^ CFU/well of *T. pyogenes* 20121 as described in the Experimental Procedure. Western blot analysis was used to detect of the levles of NLRP3 **(A)**, ASC **(B)**, Casp1(20) **(C)**, IL-1β **(D)**, IL-18 **(E)**, MMP9 **(F)**, MIF **(G)** and NF-κB **(H)** proteins; The relative protein expression levels were detected by Image J software and expressed as mean ± SD; *: *P* < .05, **: *P* < .01, ***: *P* < .001.

### *T. pyogenes* damages the respiratory tract of infected mice

To examine the effect of *T. pyogenes* infection on the development of associated pulmonary pathology in a live animal model, six-week-old C57BL/6 mice were i.p. inoculated with *T. pyogenes* or TSB, and the histopathological changes in the lungs or tracheas were examined at 1, 2, 4 and 7 dpi. The lungs of *T. pyogenes*-infected mice exhibited mild degeneration of bronchiolar epithelial cells with marked infiltration of inflammatory cells in the bronchial epithelium at 1 and 2 dpi (Fig. 4A). Furthermore, the bronchiolar epithelial cells were extensively degenerated at 4 dpi, and the adventitia showed mild edema in addition to degeneration of bronchiolar epithelial cells at 7 dpi. Compared with mock-infected mice, massive degeneration of tracheal epithelial cells was observed in mice infected with *T. pyogenes* from 1 to 4 dpi, and nuclear concentration of tracheal epithelial cells was observed at 7 dpi (Fig. 4B). Thus, *T. pyogenes* infection caused significant inflammatory response and damage in the respiratory tract of mice.

**Figure 4.**
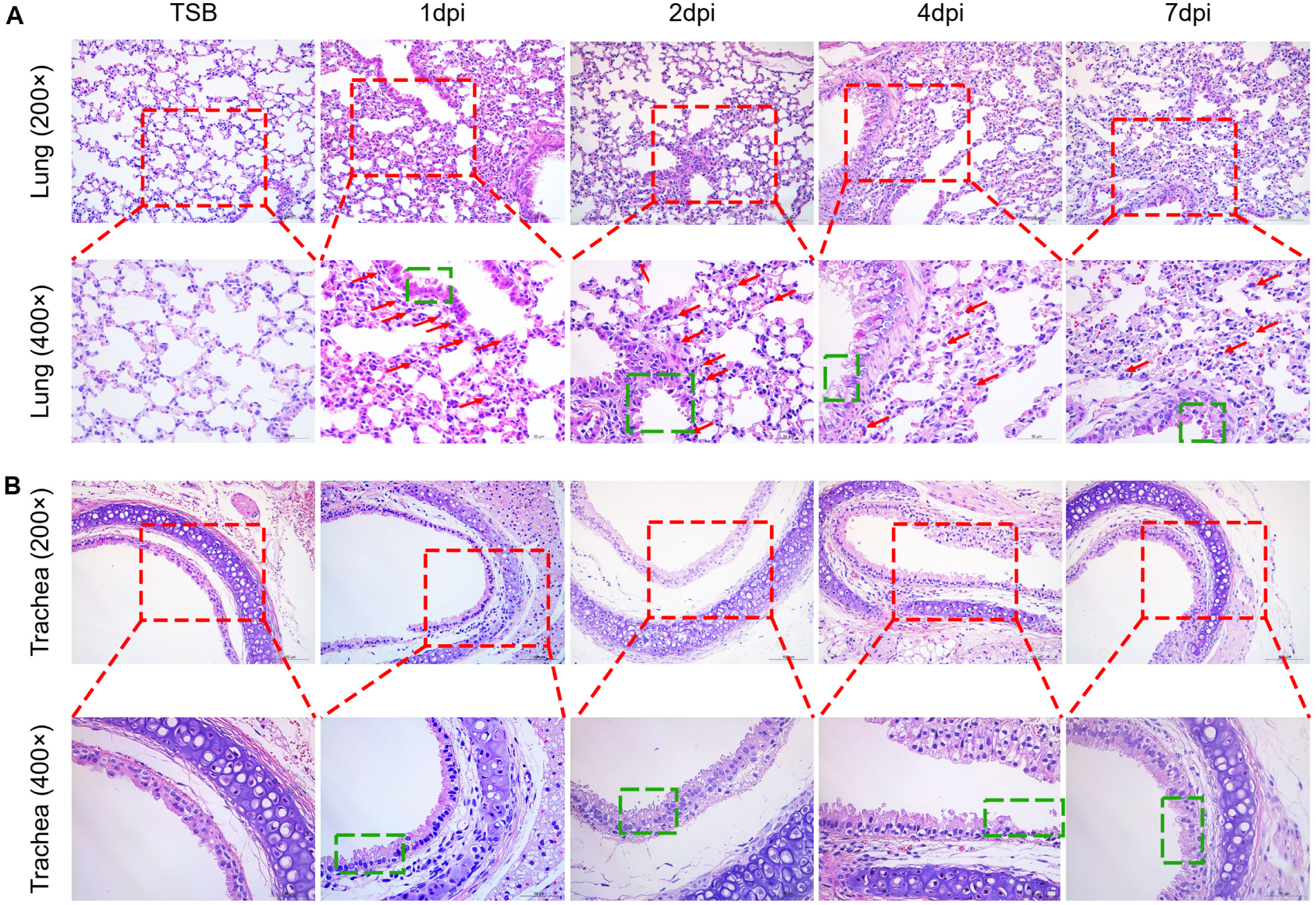
Histopathology lesions induced in the lungs of mice infected with *T. pyogenes*. Mice were infected with 2×10^6^ CFU of *T. pyogenes* strain 20121 and sacrificed at 1, 2, 4 and 7 dpi. Samples of **(A)** lung and **(B)** tracheas were fixed, embedded in paraffin and sectioned at a thickness of 5 μm. Sections were stained with hematoxylin-eosin (HE), photographed using Zeiss Viewer software; the red box shows greater detail, with extensive inflammatory cell infiltration of the lung (red arrow) and epithelial cell degeneration (green box).

### Bacterial load and inflammatory cytokines in the blood of mice infected by *T. pyogenes*

The dynamic of bacterial load in the peripheral blood of infected mice was also investigated. In agreement with our previous report (17), no bacteria were detected in the blood during the first two days of infection, while bacteria were detectable at 4 dpi (66.67 CFU/mL) and increased substantially by 7 dpi (366.67 CFU/mL) (Fig. 5A).

**Figure 5.**
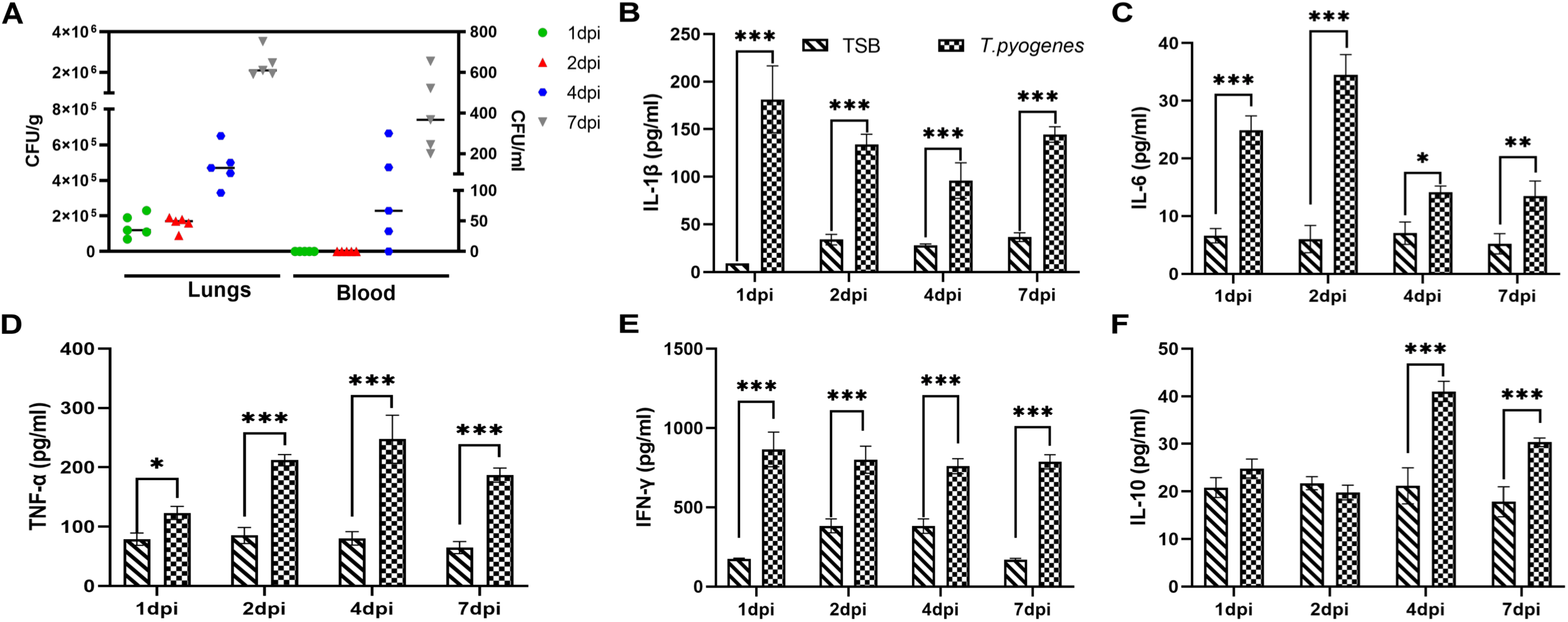
Bacterial load and cytokine levels in the lungs and blood of *T. pyogenes*-infected mice. **(A)** Bacterial load in lung (CFU/g of tissue) and blood (CFU/mL). ELISA was used to detect the levels of **(B)** IL-1β, **(C)** IL-6, **(D)** TNF-α **(E)** IFN-γ and **(F)** IL-10 in serum samples. Data are shown as the mean ± SD of the independent experiments; *: *P* < .05, **: *P* < .01, ***: *P* < .001.

However, bacterial loads in the lungs could be detected at 1 dpi and 2 dpi (1.44×10^5^ CFU/g and 1.58×10^5^ CFU/g, respectively), and bacteria levels increased strikingly at 4 dpi and 7 dpi (4.78×10^5^ CFU/g and 2.39×10^6^ CFU/g, respectively).

It is well known that airway contraction and inflammatory responses are regulated by multiple cytokines (23). To investigate the effect of *T. pyogenes* on cytokine production in mice, typical pro-and anti-inflammatory cytokines in serum were examined by ELISA kits. After *T. pyogenes* infection, IL-1β increased significantly at 1 dpi, and decreased at 2, 4 and 7 dpi compared to 1 dpi (Fig. 5B). IL-6 was increased significantly at 1 dpi (24.89 pg/mL) and peaked at 2 dpi (34.50 pg/mL) (Fig. 5C). TNF-α and IFN-γ were both significantly increased at all time-points (Fig. 5D and E), peaking at 4 dpi (247.34 pg/mL) and 1 dpi (865.29 pg/mL), respectively. IL-10 significantly increased (*P* < .001) at 4 dpi and 7 dpi (Fig. 5F).

### *T. pyogenes* activates inflammasome, MAPK, Akt and NF-κB signaling pathways in the lung of infected mice

To identify whether the inflammasome is activated in response to *T. pyogenes* infection, we used four different antibodies to detect inflammasome complexes in lung tissue by western blot (Fig. 6). The results showed that NLRP1 and NLRC4 expression (Figs. 6A and B) increased significantly between 2 and 7 dpi, while NLRP3 increased significantly only at 7 dpi (Fig. 6C) and AIM-2 was not expressed (data not shown). *T. pyogenes* clearly induced ASC expression and cleavage of pro-caspase-1, with both significantly increased from 1 dpi to 7 dpi (*P* < .001) (Fig. 6D and E). Moreover, GSDMD expression decreased significantly between 2 and 7 dpi (Fig. 6F), whereas GSDMD-N and GSDMD-C were significantly increased from 1 dpi to 7 dpi (Fig. 6G and H), suggesting pyroptosis occurred and released numerous pro-inflammatory cytokines into the infected lungs. Indeed, western blot showed a marked increase in IL-1β and IL-18 (Fig. 6I and J) during infection, MMP9 increased significantly at 1 dpi (Fig. 6K), whereas MIF had no significant changes (Fig. 6L).

**Figure 6.**
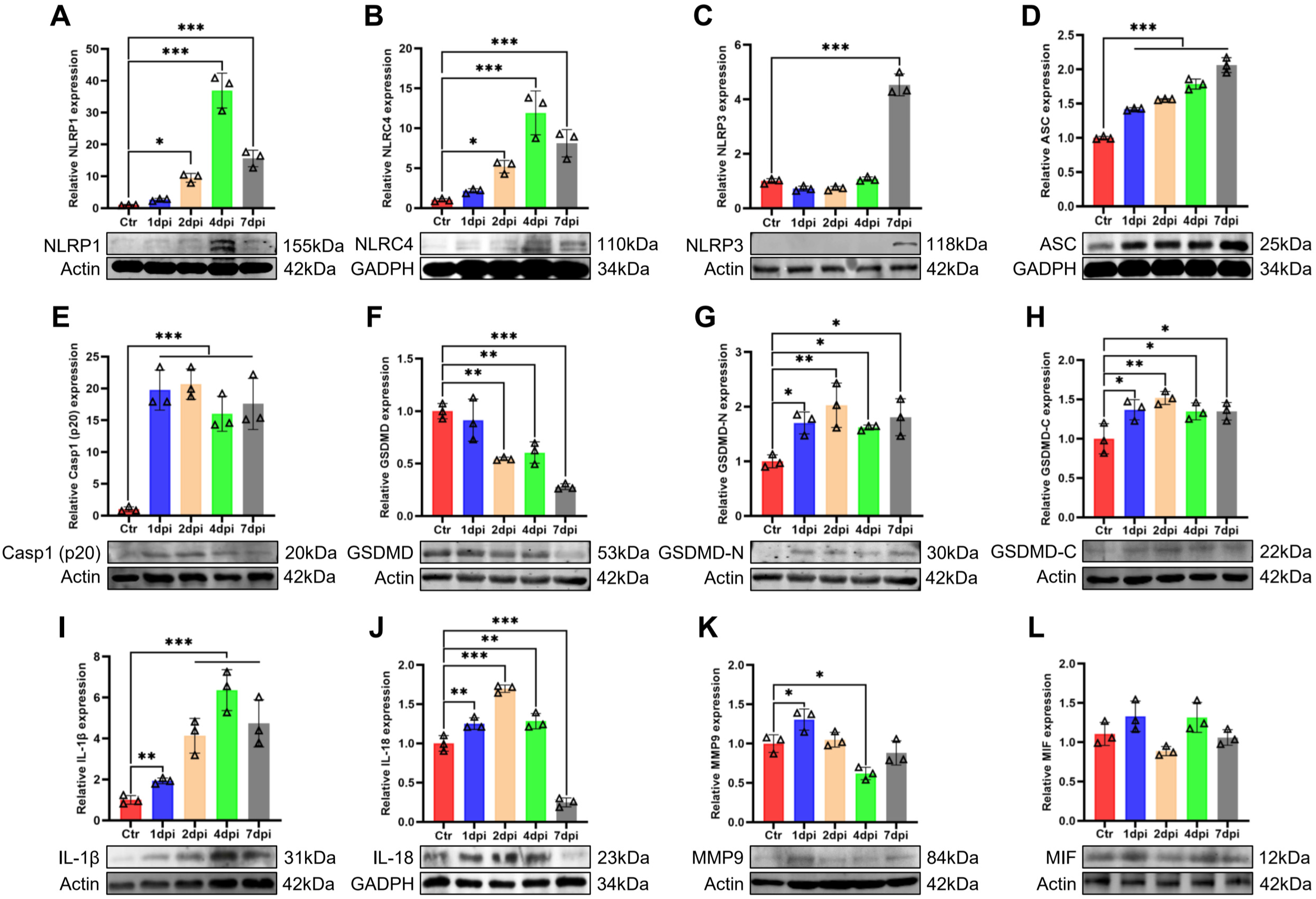
The NLR/ASC/caspase-1/IL axis is activated in the lungs of *T. pyogenes*-infected mice. Western blots of *T. pyogenes*-infected wild type mice lungs at 1, 2, 4 and 7 dpi are shown. Relative protein expression of **(A)** NLRP1, **(B)** NLRC4, **(C)** NLRP3, **(D)** ASC, **(E)** Casp1(p20), **(F)** GSDMD, **(G)** GSDMD-N, **(H)** GSDMD-C, **(I)** IL-1β, **(J)** IL-18, **(K)** MMP9 and **(L)** MIF are shown; *: *P* < .05, **: *P* < .01, ***: *P* < .001.

In order to explore the mechanism of how *T. pyogenes* may induce inflammation, we also studied the effect of infection on the MAPK signaling pathways. Western blot was used to detected classical inflammation-related signaling kinases including JNK, ERK and p38 of the MAPK pathway in lungs of infected mice. The levels of p-JNK and p-ERK were increased and the ratio of p-JNK/JNK was increased significantly from 2 dpi to 7 dpi (Fig. 7A), while the ratio of p-ERK/ERK and p-p38/p38 also increased significantly from 1 dpi to 7 dpi (Fig. 7B and C). Moreover, we tested the activation status of the canonical Akt and NF-κB pathways, which are reported to be involved in movement of immunocytes from the peripheral blood to sites of inflammation. The ratio of p-AKT/AKT was significantly increased throughout the study (*P* < .001) (Fig. 7D) and phosphorylation of NF-κB (Fig. 7E) was increased at 1, 4 and 7 dpi.

**Figure 7.**
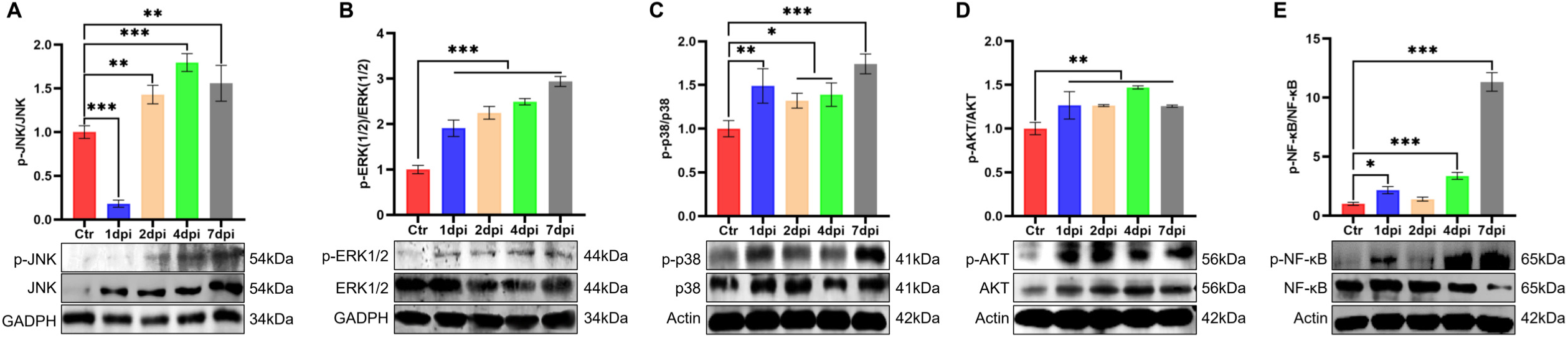
Comparison of protein expression of molecules related to activation of the MAPK pathway and inflammatory response in the lungs of mice after challenge with *T. pyogenes*. Representative western blots shows the expression and phosphorylation level of **(A)** JNK, **(B)** ERK, **(C)** p38, **(D)** AKT and **(E)** NF-κB. The mean ± SD of three experiments are shown; *: *P* < .05, **: *P* < .01, ***: *P* < .001.

### Caspase-independent and-dependent apoptosis pathways involved in lungs of mice by *T. pyogenes* infection

To determine whether *T. pyogenes* induced apoptosis in the lungs of infected mice, confocal microscopy was used to detect staining by TUNEL assay. Apoptotic signals were found not only in bronchial epithelial cells but also in the alveoli (Fig. 8A and B), which suggested *T. pyogenes* could induce apoptosis in the lungs of infected mice. Next, we used western blot to determine which type of signaling was induced by *T. pyogenes* infection. As shown in Fig. 8C, the expression of AIF protein increased significantly at 1 and 2 dpi compared with the control group. In addition, there was a significant increase in caspase-3 (Casp3) and caspase-8 (Casp8) throughout the study (Fig. 8D and E). Meanwhile, cleaved caspase-3 (c-Casp3) also increased significantly except 2 dpi and cleaved caspase-8 (c-Casp8) increased from 1 dpi to 4 dpi (Fig. 8F and G), whereas caspase-9 no significant change (Fig. 8H). These results suggest the involvement of an AIF-mediated apoptosis pathway and a caspase-dependent pathway in *T. pyogenes*-induced apoptosis.

**Figure 8.**
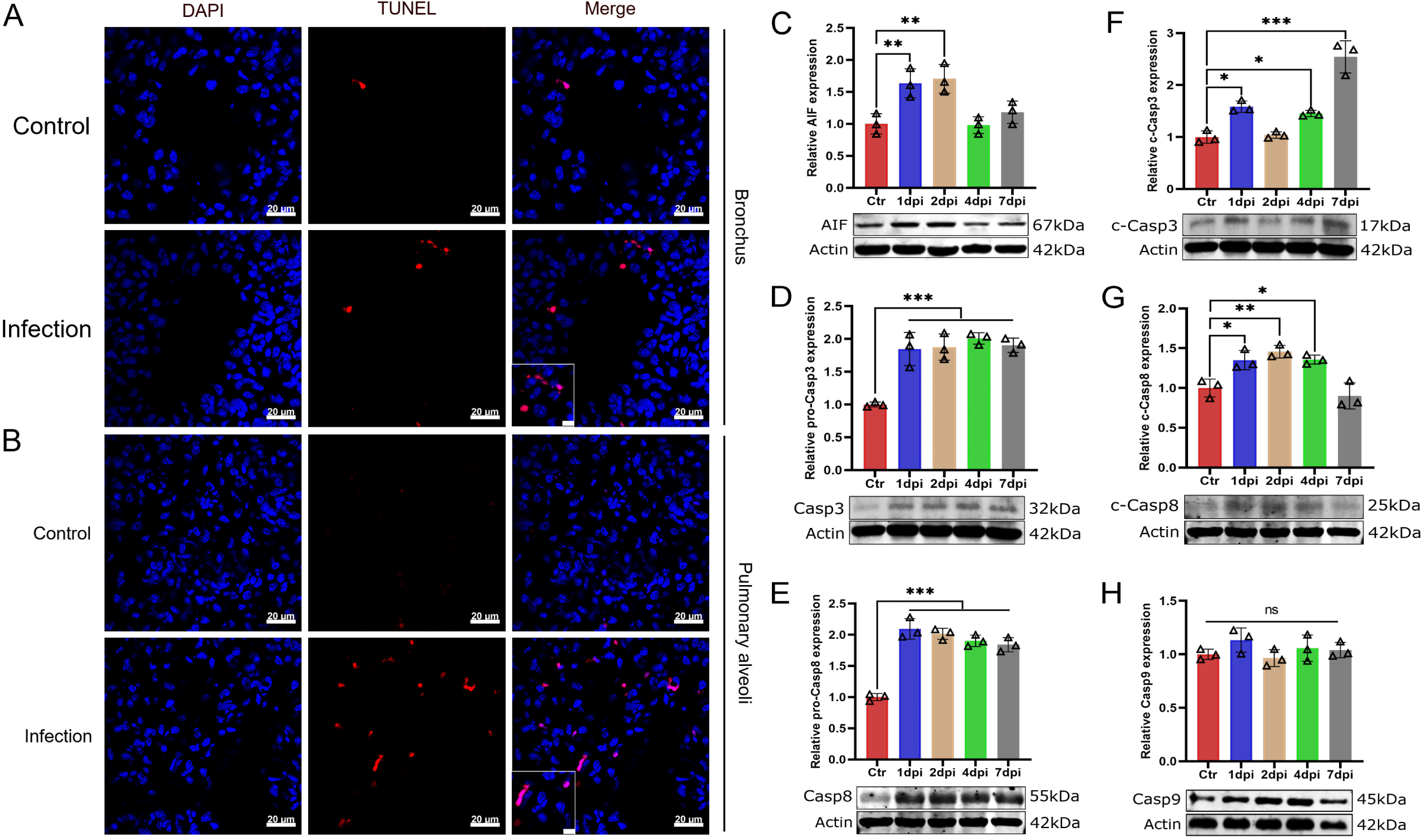
*T. pyogenes* infection induces apoptosis in mice lung cells. (A) Mice were challenged with 2×10^6^ CFU of *T. pyogenes*, and at 2 dpi, lungs were collected for immunofluorescence to verify the occurrence of apoptosis using a TUNEL assay. Apoptosis is shown red in (A) bronchial epithelial cells and (B) pulmonary alveoli in lung, and nuclei (DAPI) in blue, and the white box indicates the part view of the higher magnification images. Western blot analysis of **(C)** AIF, **(D)** caspase-3 (Casp3), **(E)** caspase-8 (Casp8), **(F)** cleaved caspase-3 (c-Casp3), **(G)** cleaved caspase-8 (c-Casp8), **(H)** caspase-9 (Casp9) expression shows apoptosis of lung cells at different days of infection. Data are shown as the mean ± SD of three experiments. *: *P* < .05, **: *P* < .01, \*\*\**P* < .001.

### TNF-α plays a role in apoptosis and inflammatory injury in the bronchiolar epithelium

The level of caspase-8 significantly increased throughout at least the first 7 days of *T. pyogenes* infection, suggesting that the TNF signaling pathway may be activated. Thus, we infected TNF-α^-/-^ mice to verify the role of TNF-α in apoptosis and lung damage. Apoptosis was detected using a TUNEL assay, and cell nuclei were stained with DAPI. As shown in Fig. 9A, there was less apoptotic signals were found in bronchial epithelial cells of TNF-α^-/-^ mice at 2 dpi, and the number of apoptotic cells in the infected TNF-α^-/-^ mice decreased significantly compared with the WT group (Fig. 9B). Furthermore, western blot analysis confirmed that the expression of caspase-8, caspase-3 and AIF decreased significantly compared with the WT group, while caspase-9 did not change significantly (Fig. 9C). Moreover, pathological sections of lung and trachea of TNF-α^-/-^ mice were observed. As shown in Fig. 9D, in the WT group, a large number of inflammatory cells infiltrated in parenchyma of lung on 2 dpi. Trachea mucosal epithelial cells partially denatured, inflammatory cells infiltrated the submucosa, and the arrangement of bronchiolar/tracheal epithelial cells was more disordered compared with TNF-α^-/-^group on 2 dpi. Whereas, in the infected TNF-α^-/-^ mice, less inflammatory cell infiltration in parenchyma of lung compared with WT group. Trachea mucosal epithelial cells showed slight degeneration and the arrangement of epithelial cells was more ordered than that seen with infection of WT mice on 2 dpi. Altogether, these data suggest that TNF-α plays a role in apoptosis and inflammatory injury to the bronchiolar epithelium of *T. pyogenes*-infected mice.

**Figure 9.**
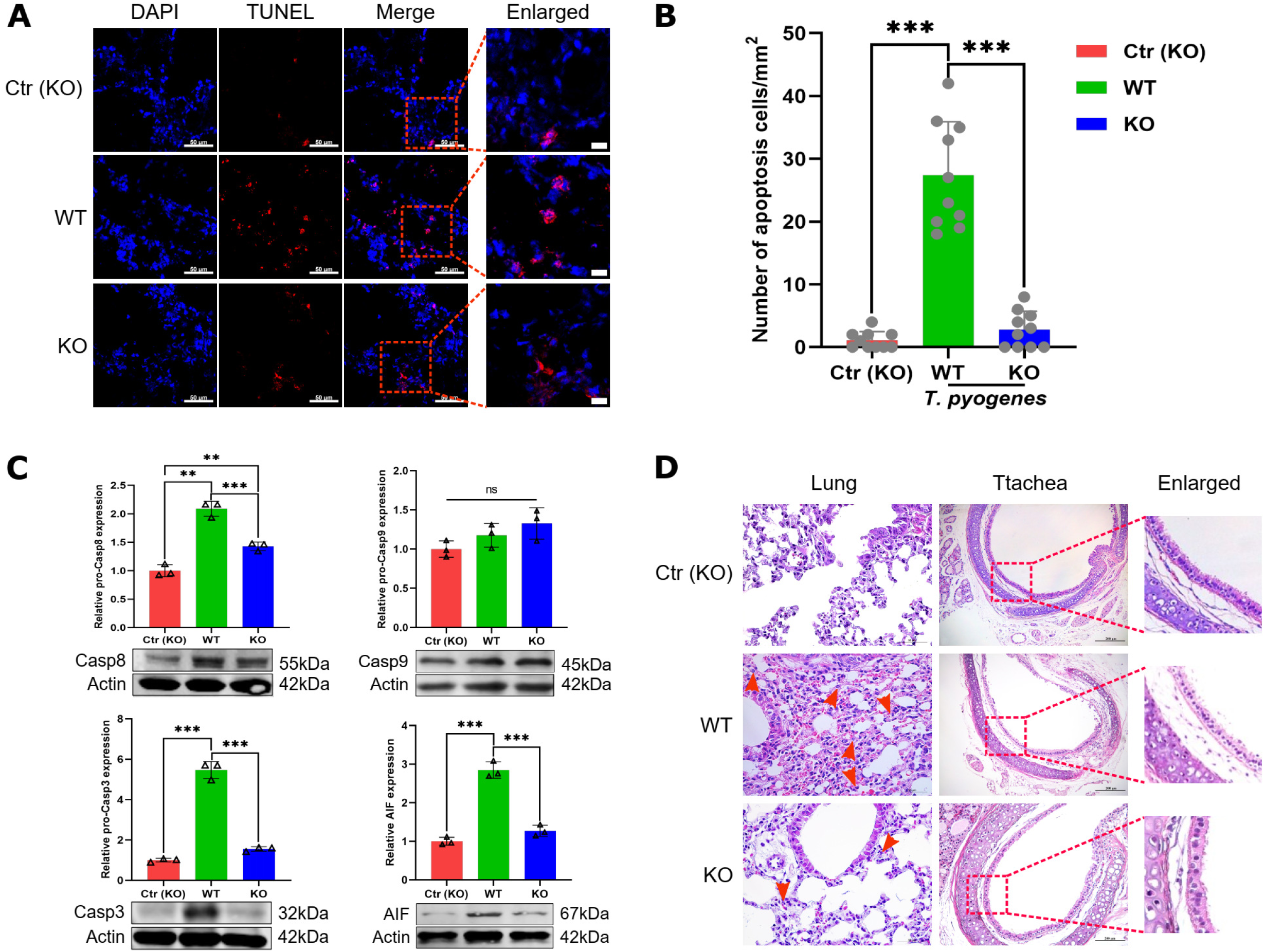
Apoptosis and injury induced by *T. pyogenes* alleviated in Lungs of TNF-α^-/-^ mice. Wild type (WT) and TNF-α^-/-^ C57 (KO) mice were challenged with 2×10^6^ CFU of *T. pyogenes*, and at 2 dpi, lungs were collected for immunofluorescence**. (A)** Apoptotic cells (TUNEL) in trachea epithelial cells were detected with the In Situ Cell Death Detection Kit, and cell nuclei were stained with DAPI. (B) Number of apoptotic cells in trachea epithelial cells from different groups of mice. \*\*\**p* < .001. Protein levels of **(C)** caspase-8, caspase-9, caspase-3 and AIF were detected by western blot. **(D)** Comparison of histopathological lesions of lungs and tracheas between infected WT and KO mice on 2 dpi, showing the epithelial layer of the respiratory tract of lungs (red dotted boxes) and neutrophil (red arrow).

## Discussion

Currently, there have less studies on the effect of *T. pyogenes* on lung functions *in vitro*. Well differentiated PBECs and PCLS make it possible to analyze the host-pathogen interaction *in vitro*, including the initial events of immune activation (24). Here we show that *T. pyogenes* is able to interact with PBECs and PCLS, where induce immune responses and develop airway damage.

In this study, notable bronchoconstriction was observed and IL-4 increase in the supernatants of infected PCLS. Since IL-4 is associated with the pathogenesis of allergic disorders in humans (25), we speculate that its increase may contribute to the bronchoconstriction, especially in the high-dose group. The mechanisms underlying IL-4 mediation of porcine bronchial constriction need to be further investigated. *T. pyogenes* could adhere to ciliated respiratory epithelial cells, growing either on surface of bronchus or within bronchoconstricted areas. Bronchoconstriction can reduce the elimination capability of ciliated cells (26). Thus, *T. pyogenes*-induced bronchoconstriction may help their proliferation within constricted areas. Furthermore, *T. pyogenes* destroyed the integrity of airway epithelium under ALI conditions, which may facilitate their movement across physical barrier of respiratory epithelium. It has been reported that the suilysin of *Streptococcus suis* can induce apoptosis of epithelial cells under ALI conditions (10). The genotype of 20121 is *plo^+^/fimA^+^/fimE^+^/nanH^+^/nanP^+^* (17). Thus, hemolysin (pyolysin, plo) may accumulate locally in *T. pyogenes*-colonized areas under ALI conditions, inducing damage to epithelial cells. This will be further verified in future studies.

The release of LDH is used to measure cytotoxicity, LDH also can be released by cells undergoing pyroptosis [36]. In this study, *T. pyogenes* induce the cytotoxicity in well-differentiated PBECs (Fig. 1A) and PCLS (Fig. 2C). According to Fig. 1A, the discontinuous cytotoxic effect may be dependent on the number of bacteria. Non-adherent bacteria were removed at 4 hpi, and those attached were too few to induce detectable cytotoxic effect at 24 hpi. Subsequently, *T. pyogenes* proliferated, reaching a threshold to manifest cytotoxicity at 48 hpi. We also test the levels of IL-1β and IL-18 in PCLS, which are released from pyrolyzed cells. Both IL-1β and IL-18 increase from 4 hpi to 72 hpi. Accordingly, the LDH increase from 4 hpi to 72 hpi, which may be contributed by pyroptosis.

*T. pyogenes* usually produces a purulent-necrotic inflammation in the lung of clinically diseased swine (5), a process regulated by cytokines. Our results confirmed that *T. pyogenes* could induce the secretions of pro-inflammatory cytokines and anti-inflammatory cytokines in PCLS and mice, which is related to the polarization of Th1 and Th2. In infected mice, pro-inflammatory serum cytokines as interleukin (IL)-1β, IL-6, TNF-a, IFN-γ are produced, which suggested type 1 T-helpers (Th1) are involved in the reaction to antigen. IL-10, a typical anti-inflammatory cytokine, was secreted in serum of *T. pyogenes* infected mice significantly increased at 4 and 7 dpi rather than at 1 and 2 dpi, which may be related to high level Th1-proinflammatory cytokine IFN-γ that inhibited Th2-polarized responses and suppress IL-10 synthesis. Our results suggested *T. pyogenes* induced a strong pro-inflammatory response at the early stage of mice infection, but anti-inflammatory process appeared relatively late. However, IL-10 significantly increased only in the 8×10^5^ CFU*-*infected PCLS group at 4 hpi rather than at 24 or 72 hpi. The time of the emergence of IL-10 is different *in vitro* and *in vivo*. The specific mechanism remains to be further studied.

In this study, *T. pyogenes* infection of PCLS and mice activated the inflammasome, which activated caspase-1, which then promoted cleavage of pro-inflammatory mediators IL-1β and IL-18 into their mature states (Fig.3). IL-1β and IL-18 are the most potent and first to be produced, which bind to their receptors to recruit and activate other inflammatory cells. IL-1β and IL-18 increased significantly after *T. pyogenes* infection of PCLS and mice, along with MMP9 and MIF. At the site of inflammation, MMP9 is often found synchronous with IL-1β (30), and MIF is considered to have a strong pro-inflammatory effect (31). This is the first report that *T. pyogenes* can induce secretion of pro-inflammatory cytokines in PCLS. Meanwhile, activation of the NF-κB pathway, which promotes the expression of pro-inflammatory cytokines including IL-6 and TNF-α in PCLS and mice (Fig.10). The NLR-ASC-caspase-1 axis and NF-κB pathway were involved in both PCLS and mice, which confirm PCLS provide a platform to analyze the early pulmonary immune response. In addition, the increased phosphorylation of JNK, ERK and AKT in mice suggested the MAPK and Akt pathways also were involved in the observed inflammation (Fig. 10).

**Figure 10.**
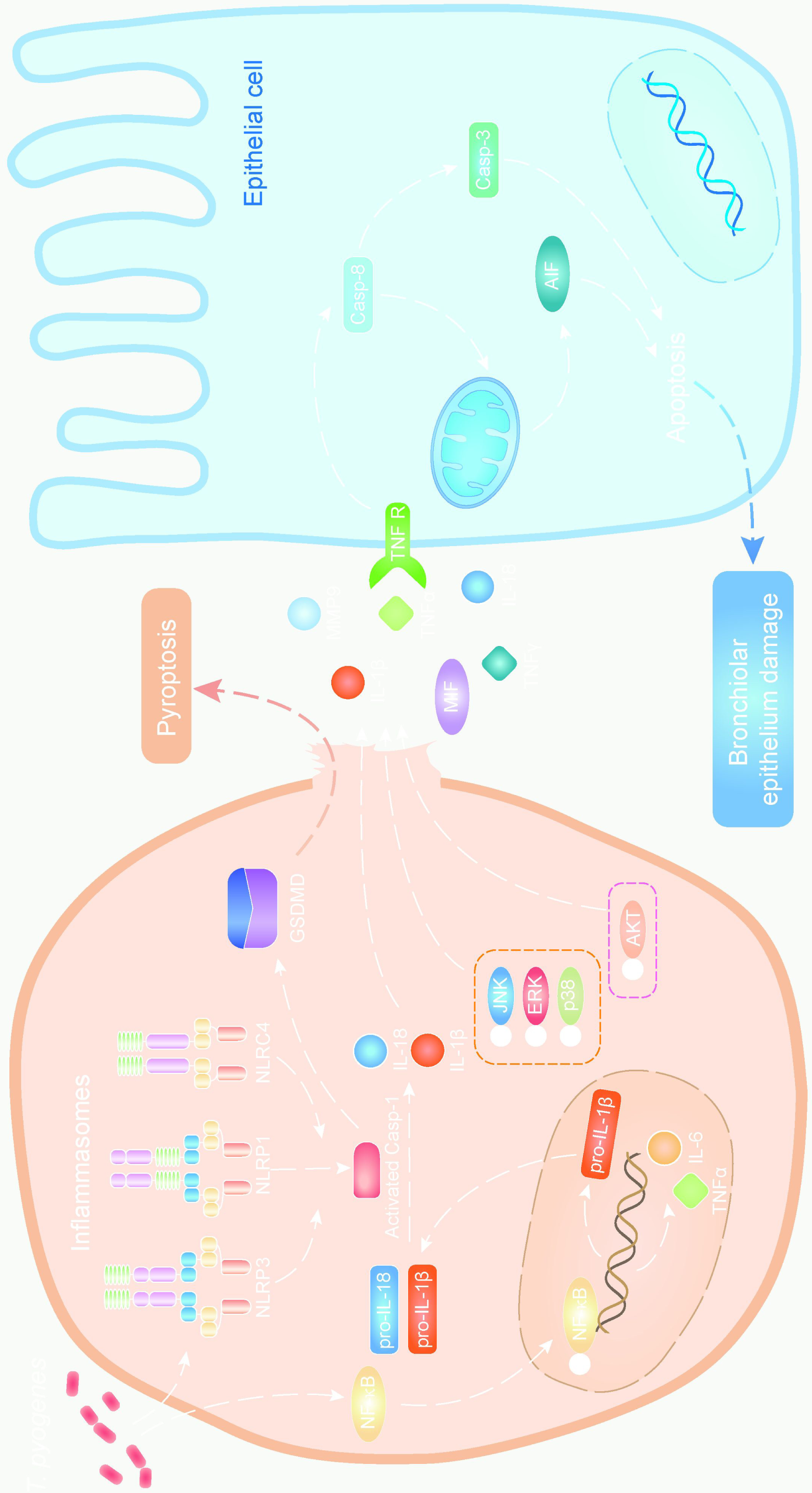
A schematic overview summarizing the main mechanism of epithelial cell injury. *T. pyogenes* infection triggers NF-κB to enter the nucleus, where it promotes expression of pro-inflammatory factors and the NLR inflammasomes. Meanwhile, *T. pyogenes* can directly promote the assembly and formation of the NLR inflammasome. Pyroptosis happens via NLR-ASC-caspase-1-GSDMD pathway, which promotes the release of mature IL-18, IL-1β and other pro-inflammatory factors into the extracellular, then cause inflammatory lung tissue injury. For example, TNF-α increases and induces epithelial cells apoptosis via TNF receptor pathway.

Since TNF-α and caspase-8 significantly upregulated in infected mice, which indicated that the TNF signaling pathway may be activated. To better understand the mechanism of leading to degeneration of epithelial cells of lungs in infected mice, we used *T. pyogenes* to infect TNF-α^-/-^ mice, found *T. pyogenes*-induced apoptosis and degeneration of epithelial cells in lungs of TNF-α^-/-^ mice were alleviated. So pro-inflammatory TNF-α plays an important role in apoptosis and lung injury induced by *T. pyogenes*. Collectively, *T. pyogenes* infection caused significant inflammatory responses in respiratory system, a variety of pro-inflammatory cytokines were produced, which are unrestricted by regulatory mediators, resulting in degeneration of epithelial cells (Fig. 10). Therefore, blocking NLR inflammasome activation to prevent large amounts of pro-inflammatory cytokines production early in infection, as well as treatments targeting TNF, may slow the disease progression in the early of infection.

In conclusion, we investigated the role of inflammation in *T. pyogenes*-induced bronchiolar epithelium damage *in vitro* and *in vivo.* NLR/ASC/caspase-1/IL axis and NF-κB pathway play the greatest role in inflammation and contribute to bronchiolar epithelium damage during *T. pyogenes* infection.

## Conclusion

The role of inflammation in *T. pyogenes*-induced bronchiolar epithelium damage are investigated *in vitro* and *in vivo*, highlighting the mechanism underlying pathological development in respiratory system during *T. pyogenes* infection.

## Author contributions

SW, FM and XC conceived the study and designed the experimental procedures. LQ, FM and SL performed the experiments. LQ, FM, HH, WZ and SW analysed the data. HH, HZ, YS, WZ, TA and XC contributed reagents and materials. LQ, FM and SW wrote the manuscript.

## Acknowledgements

This work was supported by the National Natural Science Foundation of China (32273018), the Central Public-interest Scientific Institution Basal Research Fund (NO.1610302022006) and Heilongjiang pig Modern Agricultural Technology Collaborative Innovation System.

